# A unified framework for unconstrained and constrained ordination of microbiome read count data

**DOI:** 10.1101/429340

**Authors:** Stijn Hawinkel, Frederiek-Maarten Kerckhof, Luc Bijnens, Olivier Thas

**Affiliations:** Department of Data Analysis and Mathematical Modelling, Ghent University, Belgium; Center for Microbial Ecology and Technology, Ghent University, Belgium; Quantitative Sciences, Janssen Pharmaceutical companies of Johnson and Johnson, Belgium; Center for Statistics, Hasselt University, Belgium; National Institute for Applied Statistics Research Australia (NIASRA), University of Wollongong, Australia

## Abstract

Explorative visualization techniques provide a first summary of microbiome read count datasets through dimension reduction. A plethora of dimension reduction methods exists, but many of them focus primarily on sample ordination, failing to elucidate the role of the bacterial species. Moreover, implicit but often unrealistic assumptions underlying these methods fail to account for overdispersion and differences in sequencing depth, which are two typical characteristics of sequencing data. We combine log-linear models with a dispersion estimation algorithm and flexible response function modelling into a framework for unconstrained and constrained ordination. The method allows easy filtering of technical confounders. As opposed to most existing ordination methods, the assumptions underlying the method are stated explicitly and can be verified using simple diagnostics. The combination of unconstrained and constrained ordination in the same framework is unique in the field and greatly facilitates microbiome data exploration. We illustrate the advantages of our method on simulated and real datasets, while pointing out flaws in existing methods. The algorithms for fitting and plotting are available in the R-package *RCM*.

## Introduction

Explorative visualization is a key first step in the analysis of high-dimensional ecological datasets. It provides insights into the strongest patterns in the dataset, unbiased by the researcher’s prior beliefs. It can also help to formulate new hypotheses to be tested in a subsequent study. Nowadays, microbiological communities are characterized by sequencing either marker genes or the entire metagenome of a sample, and attributing the sequences to their matching operational taxonomic units (OTUs), species or other phylogenetic levels. Throughout this paper we will refer to the lowest level to which the reads are attributed as *taxa*. Sample-specific variables, such as patient baseline characteristics or environmental conditions, can also be recorded. Microbiome sequencing datasets typically contain information on thousands of microbial taxa, whereas the number of samples and sample-specific variables is usually in the order of tens to hundreds. These data are thus *high-dimensional*, and require a dimension reduction before visualization. Apart from the biological variability, the DNA-extraction, amplification and sequencing steps, introduce additional variability and technical artefacts, such as differences in sequencing depth. At best, data visualization methods must be insensitive to this technical noise, while accurately capturing the biological signal. The first aim of such a dimension reduction is to optimally represent (dis)similarities between samples in an *ordination*: samples that are similar in high dimensional space should also be represented close together in a two or three dimensional visualization. A second aim is to elucidate which taxa drive the (dis)similarities between samples. A final objective might be to identify which sample-specific variables can explain the (dis)similarities in taxa composition between samples. Current ecological ordination methods mainly focus on the first aim, which is to ordinate samples optimally in few dimensions. These approaches often fail to elucidate which bacterial taxa differentiate the samples. Moreover, methods that attempt to visualize variability in a dataset (unconstrained ordination) and methods that explore the role of sample-specific variables in shaping the community (constrained ordination), have evolved independently.

A very popular ordination method for the microbiome is *principal coordinates analysis* (PCoA) [1], also known as *multidimensional scaling* [2]. First, the data analyst chooses a particular distance measure, which is calculated for every pair of samples in the high-dimensional space. Next, samples are represented in two dimensions such that their pairwise Euclidean distances approximate their corresponding distances in high dimensional space as close as possible. However, no matter how well motivated the choice of distance measure for a particular application, the contribution of the individual taxa to the separation between the samples is lost in the distance calculation (taxon scores are sometimes added to the PCoA plots as weighted sample scores, but they do not reflect their contributions to the distance measures); see Fig 1A. Moreover, distance-based approaches have been shown to be affected by differences in dispersion [3] and library sizes [4, 5] between the samples.

**Fig 1.**
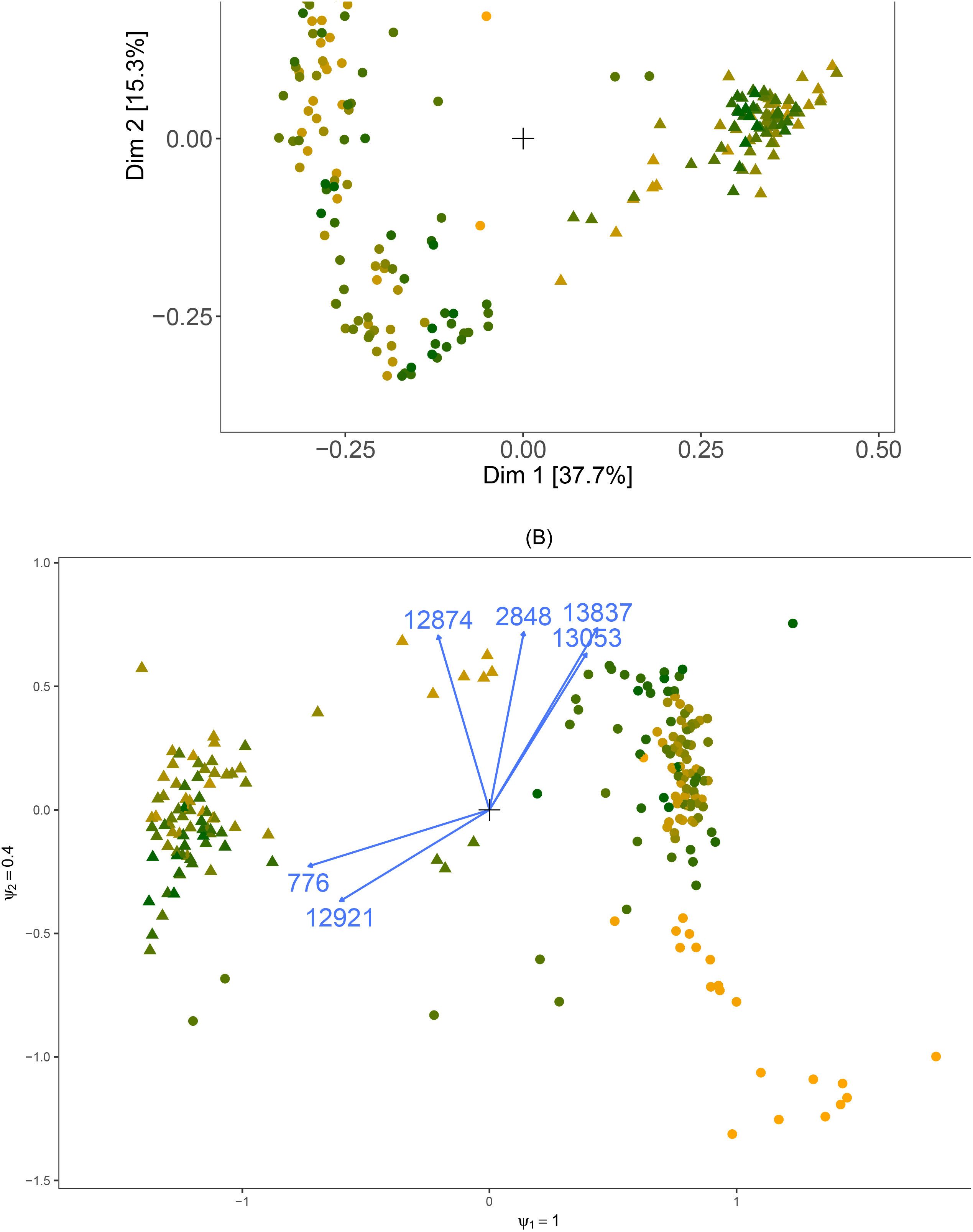
Unconstrained ordination methods. (A): Principal coordinates (PCoA) sample ordination with Bray-Curtis distances on relative abundances of the Turnbaugh mice dataset. Dots represent mice, percentages on the axes indicate fraction of eigenvalue to the sum of all eigenvalues. (B): Biplot of the unconstrained RC(M) ordination of the same dataset. Arrows represent taxa, the ratios of the parameters reflect the relative importance of the corresponding dimensions. Only the six taxa with strongest departure from homogeneity are shown for clarity. The sample ordination is similar to PCoA, but the RC(M) also identifies which taxa contribute most to the separation of the samples. LF/PP: low fat, plantpolysaccharide rich.

Correspondence analysis (CA) [6] is a classical statistical method for the exploration of contingency tables, which allows for quantification of taxon contributions to the sample ordination. Canonical correspondence analysis (CCA) [7] even allows to restrict the sample ordination to be explained by sample-specific variables (see Fig 2A). This technique thus allows for unconstrained (CA) and constrained (CCA) analysis in the same framework, which greatly enhances their use for researchers. Correspondence analysis relies on residuals for capturing the discrepancy between observed counts and the counts expected in case of identical taxa composition in all samples (sample homogeneity). It implicitly assumes a certain mean-variance relationship for normalization of these residuals. However, a residual-based approach is not well adapted to skewed data, and its mean-variance assumption is too rigid to account for the overdispersion which is typically encountered in sequencing data [3]. Moreover, both CA and CCA implicitly assume unimodal response functions, i.e. for each taxon the expected abundance shows a bell-shaped functional relationship with a *score*. This score may be latent (CA) or observed (CCA), and represents the value of a particular sample along a *gradient* of e.g. environmental conditions. CCA makes strong assumptions on the shape of these taxon response functions [7, 8].

**Fig 2.**
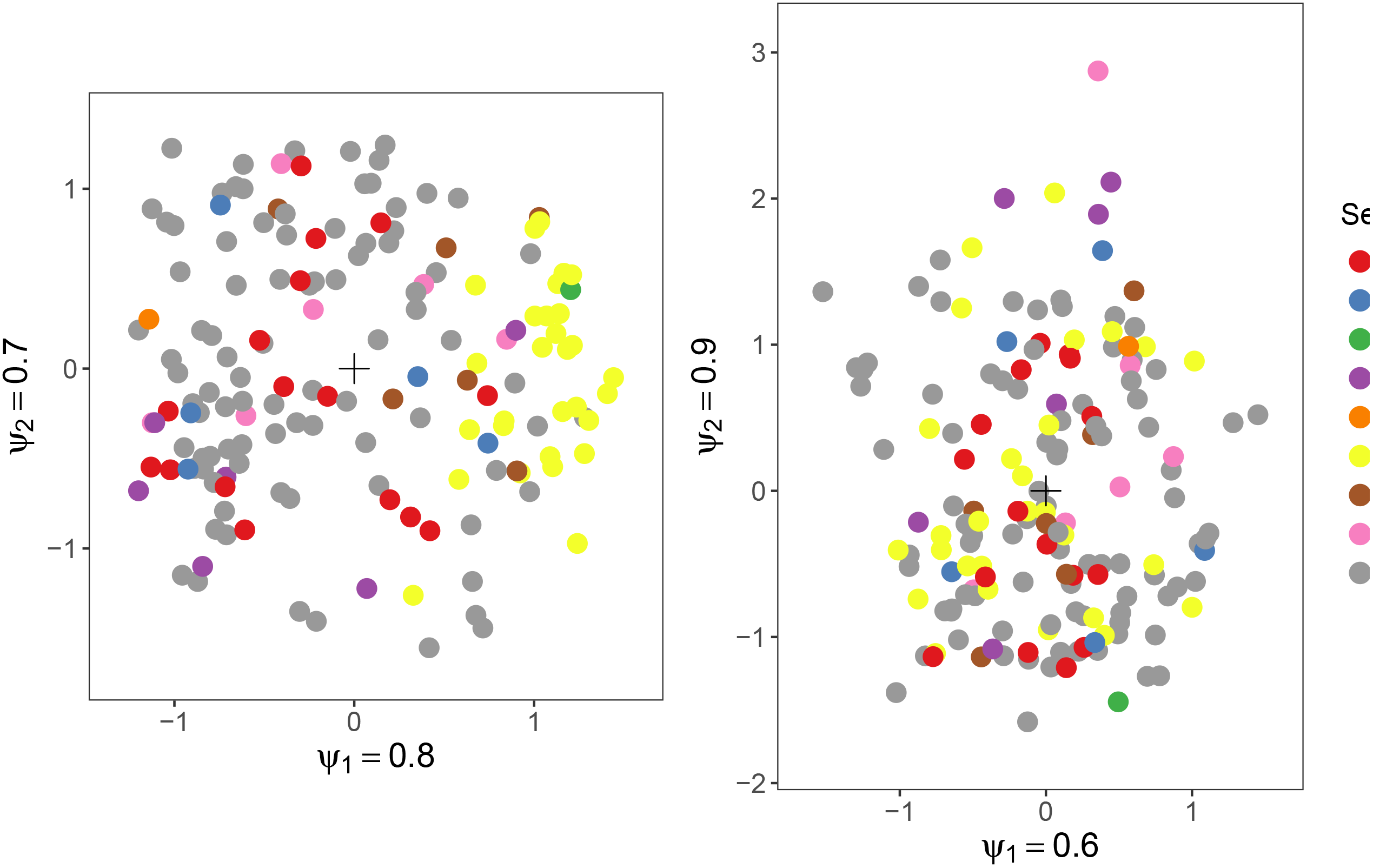
Constrained ordination methods. (A): Triplot of canonical correspondence analysis (CCA) of the Zeller data. Dots represent samples, the taxon labels indicate the location of the peaks of the taxon response functions under strict assumptions. For clarity, only the eight taxa with peaks furthest from the origin are shown. Percentages along the axes indicate fractions of total inertia explained by the dimension. Arrows depict the contribution of the variables to the environmental gradient. (B): Triplot of the constrained ordination of the same dataset by the RC(M) method with linear response functions. Arrows represent taxon response functions, and labels represent variables constituting the environmental gradient. The ratio of the ψ parameters reflects the relative importance of the corresponding dimensions. Only the eight taxa that react most strongly to the environmental gradients (the longest arrows) are shown. Two Fusobacterium species are among the taxa most sensitive to the environmental gradient, and are more abundant in cancer patients than in the others, which is in accordance with the findings of [9].

Recently, new data visualization methods for sequence count data have been proposed that aim to account for their compositionality [10]. Compositional data are constrained to a constant sum that is unrelated to their composition (e.g. the library size for sequencing data). As a result, only the proportions of the components (e.g. taxa) are meaningful, and an increase in proportion (relative abundance) of one taxon automatically entails a decrease in proportion of some other taxon or taxa. These visualization methods take the compositional nature of the data into account by working on log-ratios of relative abundances, and allow to visualize the role of the taxa in the ordination. However, since sequencing count tables have very high zero count frequencies, working with log-ratios requires the addition of pseudocounts before log-transformation to avoid division by zero. The choice of the pseudocount is arbitrary and can strongly affect the eventual ordination [11]. In addition, normalizing to relative abundances and using ratios, discards the information in the library size and taxon abundance, and associated variance. As a result these methods fail to account for heteroscedasticity, and can be distorted by technical artefacts such as differences in library size).

Row-column interaction models have been used previously for ordination [12, 13], but only for unconstrained ordination. Moreover, some assume inappropriate distributions with a common dispersion parameter for all taxa [12] or do not treat samples and taxa on an equal footing [13].

As the preceding examples illustrate, a rich literature exists on ordination of ecological data, but few methods bridge the gap between unconstrained and constrained ordination. Correspondence analysis [6, 7] is a rare exception, but it is too restrictive for sequencing count data. Other methods have no counterpart for constrained analysis [10, 12, 13], or resort to inefficient two-step approaches [14]. On the other hand, many methods for constrained ordination focus solely on the estimation of either the gradient or the response curve. As a result, they do not produce comprehensive triplots which simultaneously show the relationships between samples, taxa and sample-specific variables [8, 15, 16].

Upon combining ideas of log-linear analysis of contingency tables [17, 18], dispersion estimation for sequencing data [19] and flexible response function estimation [8, 20], we propose a new row-column interaction model for the visualization of the strongest signals in a microbiome count dataset. As it is based on a statistical regression model, our approach has the flexibility to correct for known confounders such as sequencing center or technology, and to adequately deal with the mean-variance relationships of sequencing data. Our method integrates unconstrained and constrained ordination into the same framework, will simplifies the workflow of a microbiome data exploration. It is implemented in R [21] in the form of the *RCM* package, which enables the creation of annotated graphs of the ordinations. Unlike many other ordination methods, the underlying assumptions of our method are explicitly stated and can be verified through simple diagnostic plots.

Comparisons of ordination methods have mainly focused on sample ordination, either from the viewpoint of ordination along a gradient [3, 22–26] or clustering [4, 27], and have failed to identify a single best method. They rely mainly on simulated data based on gradients with hypothesized response functions [22–25, 28], and on clusters of samples with similar compositions [3, 25, 28] or on real datasets with supposedly known gradients or clusters [3, 25, 26, 28, 29]. Few studies pay attention to the role of the taxa in the ordination, but none of them does so in a quantitative way [3, 28, 30, 31]. Here we present a simulation study that evaluates sample ordination as well as identification of taxa that contribute to the separation of the samples.

## Materials and methods

Computations were run on a Dell laptop, on two servers with 12 respectively 30 cores and on the high performance computing facilities of VSC (the Flemish Supercomputer Center). All analyses were run with the R programming language versions 3.4.3 and 3.3.1 [21]. All R-code used for the publication is available in the S1 File. The code for fitting and plotting the RC(M) models can be found in the R-package *RCM*, which can be installed from https://github.com/CenterForStatistics-UGent/RCM.

### Datasets

The Human Microbiome Project (HMP, V13 region of the 16S rRNA gene) [32] and the American Gut Project (AGP) [33] provide microbiome count datasets of healthy human volunteers. Data from two studies on the colorectal microbiome of cancer patients, referred to as the Zeller data [9] and the Kostic data [34] are also included. Furthermore, a study on several generations of gnotobiotic mice, referred to as the Turnbaugh data [35], provides non-human microbiome data. A study on microbes in cooling water provides data from a non-mammalian source, referred to as the Props data [36]. All datasets are available in the S2 File.

### Simulation study

Simulations were set up by assuming a particular count distribution, for which the parameters were estimated from a real dataset. Parameter values for the taxa and samples were then sampled from this pool of realistic parameter estimates for every Monte Carlo simulation. We chose the negative binomial, zero-inflated negative binomial and Dirichlet multinomial as count distributions. The Dirichlet multinomial distribution generates much higher zero frequencies than observed in microbiome data, but it was included because of its common use in microbiome science [37]. Parameter values were obtained as follows. Library sizes were randomly sampled from a pool of observed library sizes of the HMP datasets. The taxon-wise mean abundance and dispersion parameters from the negative binomial distribution were estimated by maximum likelihood from the mid vagina, stool and tongue dorsum samples from the HMP and from the AGP data. The overdispersion parameter of the Dirichlet multinomial was estimated from the AGP dataset using the method of moments. The mixing proportions of the zero-inflated negative binomial were estimated by maximum likelihood from the AGP data. Datasets were generated with 60 samples and 1000 taxa.

Two sets of scenarios were considered. In a first set no biological signal was introduced. The first scenario consisted in simulating data with the negative binomial distribution such that in each of four groups of 15 samples, the sampled library sizes were multiplied with a constant: 0.2, 1, 5 and 10 for the four groups. This generates technical variability that should not be picked up by the ordination method. The second scenario was similar, but now the sampled taxon-wise dispersions were multiplied by 0.2, 1, 2 and 5 for the four groups. The second set of scenarios were designed to represent different types of biological signal that should be detected and visualized by the ordination method. Counts were also generated for 4 equally sized groups of samples, but with different taxa compositions.

In the first scenario, which will be referred to as NB, initially one taxa composition was sampled for all the groups. This composition was then altered for every group separately by multiplying a random sample of 10% of the taxon abundances by a fold change of 5 so as to make them differentially abundant (DA). Counts were generated with the negative binomial distribution. The second setting, referred to as NB(cor), was identical to the first, except that counts were generated with between-taxon correlations. These taxon correlation networks were estimated by SpiecEasi [38] on the mid vagina, stool and tongue dorsum datasets of the HMP and on the AGP data. A correlation network was sampled for every Monte Carlo instance. The third scenario, referred to as NB(phy), was also similar to NB, only now a random phylogenetic tree was created for every dataset. Next, the tree was divided into 20 clusters of related taxa, and differential abundance was introduced in one of the clusters with a fold change of 5. This assures that the DA taxa are phylogenetically related, similar to the second approach in [39]. The fourth simulation scenario, which will be referred to as DM, used the Dirichlet multinomial distribution, for which DA is introduced as for the NB scenario. The fifth scenario, referred to as ZINB, was again similar to the NB setup, but used the zero-inflated negative binomial distribution. The DA is introduced only in the count part of the distribution. Further details and additional simulation scenarios can be found in Section 3.1 of the S1 Appendix.

### Competitor ordination methods

As competitor ordination methods we include (1) detrended correspondence analysis (DCA), (2) ordination through PCoA with (a) Bray-Curtis dissimilarities on absolute abundances (Bray-Curtis-Abs), rarefied absolute abundances (Bray-Curtis-rare), relative abundances (Bray-Curtis) and log-transformed abundances (after adding a pseudocount of 1) (Bray-Curtis-Log), with (b) Jensen-Shannon divergence (JSD), with (c) unweighted and weighted UniFrac distances (UniFrac and w-UniFrac), and (3) ordination through non-metric multidimensional scaling with Bray-Curtis dissimilarities on relative abundances (Bray-Curtis-NMDS) and (4) DPCoA using *phyloseq* [40]. Correspondence analysis approximating the Pearson’s chi-squared (CApearson), the contingency ratio (CAcontRat) and the chi-squared distance (CAchisq) was implemented according to [41]. The ordination based on the Hellinger distance (Hellinger) follows [42]. Compositional data analysis (CoDa) biplots follow [10]. The *gomms* R-package was used to run the GOMMS ordination method [12] and the *tsne*R-package for the t-SNE method [43]. All methods were applied to count matrices trimmed for taxa below a prevalence threshold of 5% or with total count lower than 10% of the number of samples. Ordinations in three dimensions were requested.

### Performance metrics

The results of all ordinations on the simulated datasets were evaluated for separation of the sample clusters through silhouettes [44] and through a pseudo F-statistic [29, 45]. The contribution of the taxa to the correct separation of the samples is quantified by the “taxon ratio”. This metric is based on the average inner product of the DA taxon scores and the samples scores of samples in which the taxa are known to be differentially abundant. This yields a measure of how much these DA taxa contribute to the separation of the samples. The mean inner product of the non-DA taxon scores with the same sample scores should be small for an ordination method that can discriminate between DA and non-DA taxa. The ratio of the former to the latter mean inner product is the taxon ratio. It is used as a measure of method performance in terms of taxon identification. Finally, also the correlations of the sample scores with the observed library size are calculated. These summary measures allow a quick evaluation of all simulation runs, but inevitably high values for these measures do not always correspond to an aesthetically pleasing biplot.

## Results

### The RC(M) model

#### The unconstrained RC(M) method and biplots

A typical microbiome count dataset is represented as an n × p count table **X** for *n* samples and *p* taxa. An n × d matrix of sample-specific variables **Q** (the metadata) can also be available; categorical variables are represented by 0*/*1 dummy variables. In the unconstrained RC(M) model, the expected count of taxon *j* in sample *i* is modelled as 
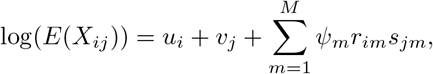
in which *u_i_* + *v_j_* represents the independence model. The independence model describes the expected counts under the assumption of equal taxa composition in all samples (sample homogeneity). In the current context, exp(*u_i_*) is a measure of sequencing depth of sample *i*, and exp(*v_j_*) is the mean relative abundance of taxon *j*. The factor *r_im_* is a sample score that captures departure from homogeneity in sample *i* in dimension *m*, and *s_jm_* is a taxon score for taxon *j* in dimension *m*. Because the sample and taxon scores are normalized (see Section 2.1.5 of the S1 Appendix), the parameter ψ*_m_* is a measure of overall strength of departure from homogeneity in dimension *m*. The constant *M* is the number of dimensions of the ordination, which is usually 2 or 3. This mean model is augmented with a negative binomial count distribution for *X_ij_*, which captures the high variance and high zero frequency in microbiome count data [3, 27]. The term in 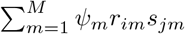 Equation 1 can be used to make interpretable biplots for visualizing departures from homogeneity. In 2D one can plot ψ_1_*r_i_*_1_ versus ψ_2_*r_i_*_2_ to obtain a sample ordination plot. Samples close together on this plot depart similarly from homogeneity and thus have similar taxa compositions (see Fig 1B). To reveal the role of the individual taxa in this ordination, we add the *p* taxon scores *s_j_*_1_ versus *s_j_*_2_ as arrows on the same plot. The orthogonal projection of (*s_j_*_1_*, s_j_*_2_) on (ψ_1_*r_i_*_1_, ψ_2_*r_i_*_2_) gives 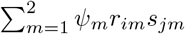, which quantifies the deviation of taxon *j* in sample *i* from sample homogeneity; see Equation 1.

Loosely speaking, taxa have a higher expected abundance in samples for which the sample dots and taxon arrows lie at the same side of the origin, and a lower expected abundance if they lie at opposite sides. See Section 2 of the S1 Appendix for a detailed description of the estimating algorithm and the construction of biplots, Section 4 for real data examples.

#### Conditioning in the RC(M)-model

Technical sample-specific variables such as sequencing center and technology are known to affect the observed counts [46]. When these confounding variables are known, they can be included in the RC(M) model. This effectively filters out their effect, similar to conditioning in correspondence analysis [47]. With **G** an n × k confounder matrix (a subset of **Q**), model 1 is extended to 
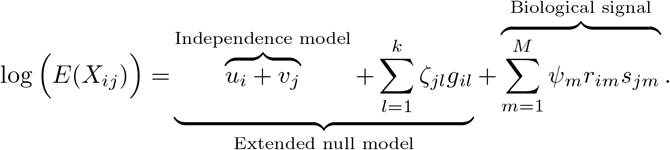

In this model, ζ*_jl_* is a parameter such that the interaction term ζ*_jl_g_il_* captures the departure from homogeneity of taxon *j* in sample *i* due to variable *l*. As a result, the biological signal term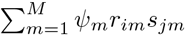 is free of the effect of the confounding variables. This is illustrated in Fig 3. Details can be found in Section 2.1.4 of the S1 Appendix. Conditioning on known confounders can be applied in the unconstrained as well as in the constrained RC(M) model (see next section).

**Fig 3.**
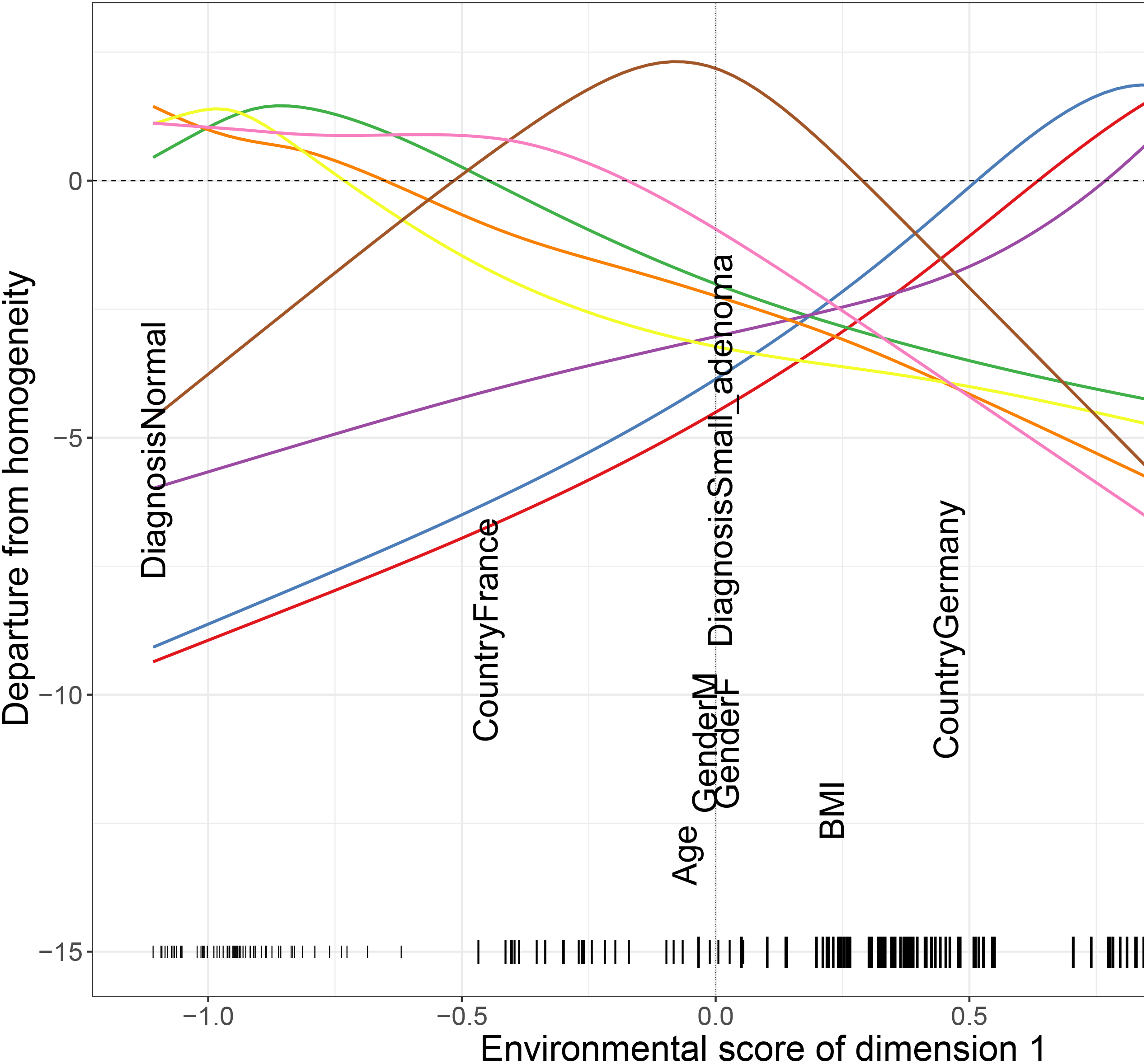
Effect of conditioning on unconstrained RC(M) ordination. Left: Unconstrained RC(M) sample ordination of the anterior nares samples of the HMP dataset without conditioning. Right: Ordination of the same sample, but after conditioning on the main sequencing center (Washington University genome center (WUGC), J. Craig Venter Institute (JCVI), Baylor College of Medicine (BCM) and Broad Institute (BI)). The ratio of the ψ parameters reflects the relative importance of the corresponding dimensions.

#### The constrained RC(M) model

The idea of a constrained ordination is to visualize the variability in the dataset that can be explained by sample-specific variables [7, 8]. Constrained ordination is traditionally performed by finding an environmental gradient *_m_* for every dimension *m*. Let **c**_*i*_ represent the *i^th^* row of **C** (a subset of **Q**, excluding **G**) containing the sample-specific variables for which one wishes to investigate the effect on the taxa composition. The environmental gradient then defines an environmental score 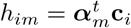 for every sample *i*. This *h_im_* can be seen as an equivalent of the row score *r_im_*, but constrained to be a linear combination of sample-specific variables. Each taxon *j* is allowed to react to this environmental score in a different way through taxon-specific response functions *f_jm_*(*h_im_*). The constrained RC(M) model then becomes 
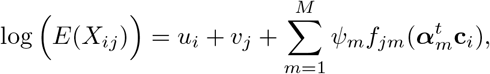
in which *u_i_, v_j_* and ψ*_m_* play the same role as in models 1 and 2. The difference with the classical gradient analysis methods is that we use the response functions to model the *departure from homogeneity*. In this way our method automatically accounts for differences in sequencing depth and taxon abundance. The environmental gradient α*_m_* is estimated by maximizing the likelihood ratio between a model with the taxon-specific response functions *f_jm_* of model 3, and a model with a common response function, *f_m_* = *f*_1*m*_ = *f*_2*m*_ = *…* = *f_pm_*, for all taxa. This encourages maximal niche separation between the taxa [8]. The correct shape of the response function has been the subject of theoretical debate [15, 16, 48], but it evidently depends on the taxon, as well as on the available sample-specific variables and their observed values. For easy interpretability we propose to use linear response functions *f_jm_*(*h_im_*) = β_0*jm*_ + β_1*jm*_*h_im_*, analogous to redundancy analysis [49]. These response functions can easily be represented in two dimensions by an arrow originating in 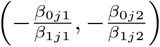, with 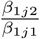 and magnitude proportional to 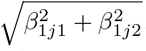

**Fig 4.**
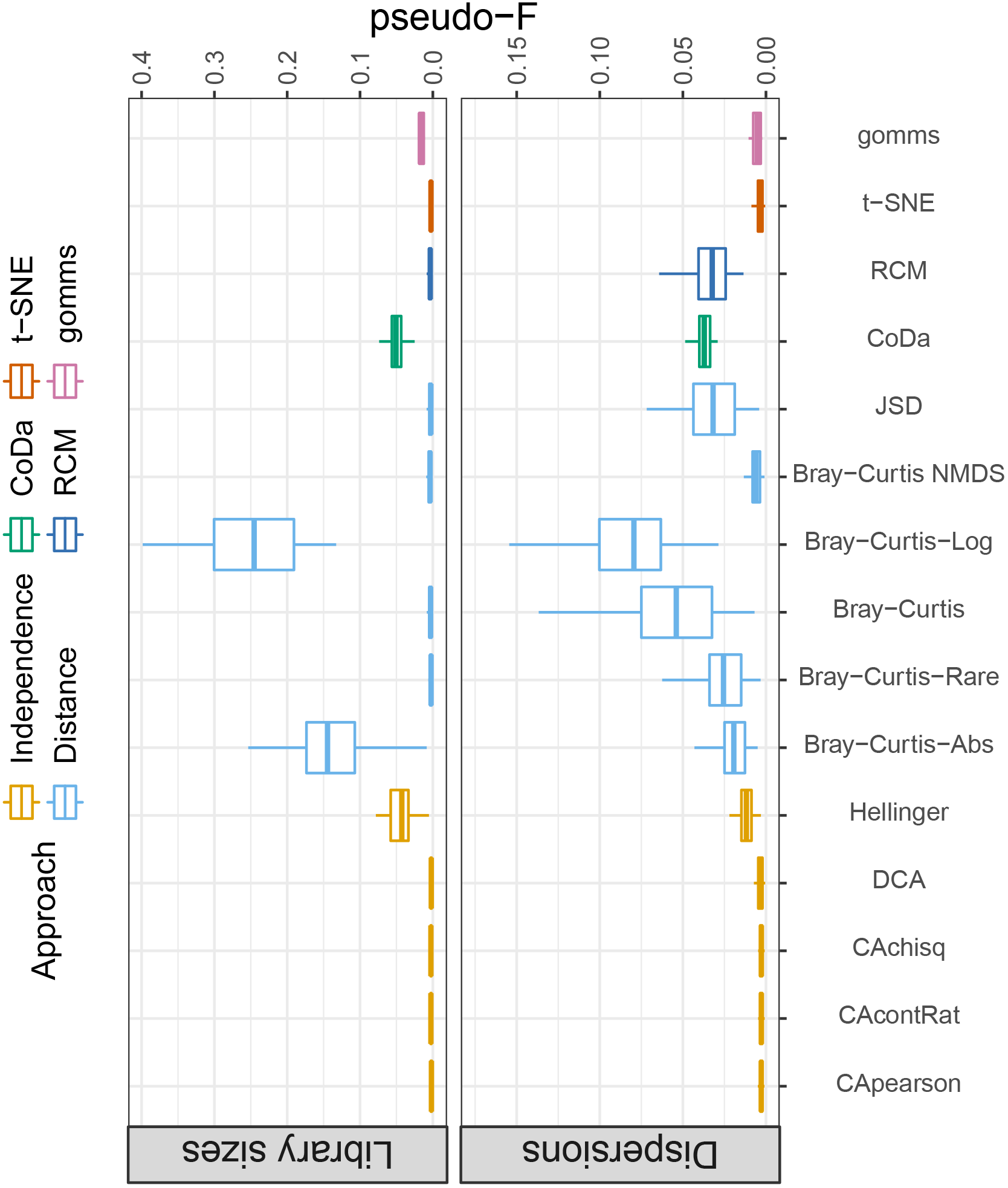
RC(M) ordination with nonparametric response functions. One-dimensional triplot of the first dimension of the constrained RC(M) ordination with non-parametrically estimated response functions of the Zeller data. Coloured lines represent taxon response functions. The horizontal dotted line represents the expected taxon abundances under sample homogeneity. Only the eight taxa that react most strongly to changes in the environmental score are shown for clarity. Black labels show the variables constituting the gradient and vertical dashes at the bottom represent the sample scores. The horizontal positions of the variable labels indicate how much they contribute to the environmental gradient; the vertical stacking is only for readability.

A more flexible approach to modelling the taxa niches is provided by non-parametric estimation of the response functions with generalized additive models (GAMs) [50], similar to [20]. It provides possibly improved constrained sample ordination and gradient estimation, but also allows the researcher to study the way the taxa react to the environment with less prejudice. Fig 4 shows that different taxa can react entirely differently (and non-linearly) to changes in their environment. Quadratic response functions are frequently used implicitly [7] or explicitly [8, 51] to model unimodal response functions; they are also implemented in the RCM R-package. They are, however, harder to depict in a triplot than linear response functions, while still providing less flexibility than non-parametrically estimated response functions. Moreover, for some taxa the estimated parameters of quadratic response functions may make the response curve convex rather than concave [52].

#### Diagnostic tools for the RC(M) ordination

Almost all ordination methods come with certain assumptions, but they are rarely explicitly mentioned, let alone checked by the user ([13] is a notable exception). The likelihood framework by which the RC(M) model is fitted, explicitly states model assumptions, and allows these to be checked. Deviance residuals are a standard diagnostic tool in generalized linear models [53], and can be used to detect taxa and samples that poorly fit the model. Influence functions can help to identify samples or taxa with a dominant role in shaping the final ordination [54]. Both of these diagnostic plots can point researchers to outlying and possibly interesting samples and taxa that deserve further scrutiny (see Section 2.4 of the S1 Appendix for examples).

**Fig 5.**
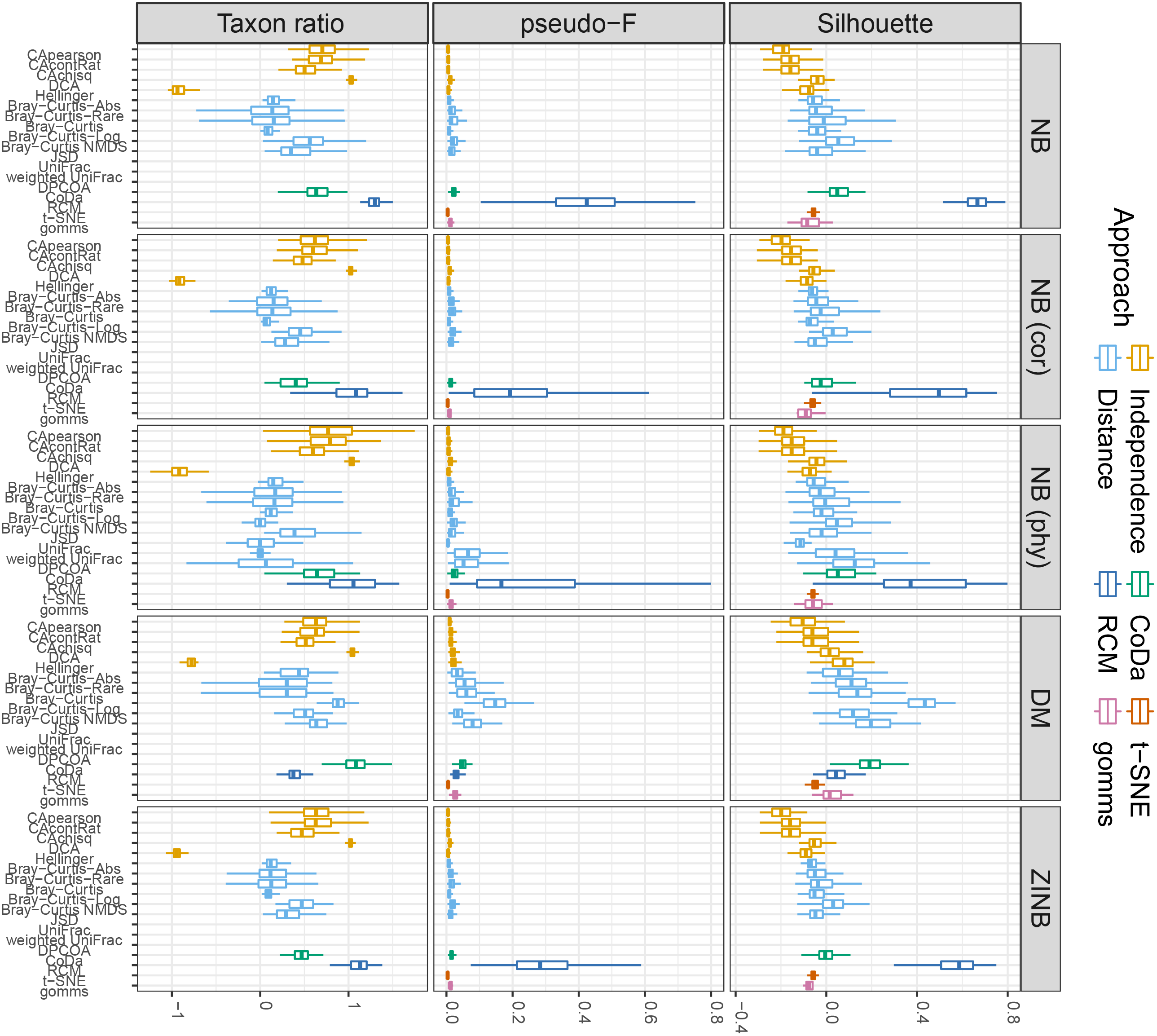
Results of simulations without signal. Boxplots of the pseudo-F statistic for sample clustering (y-axis) for several ordination methods (x-axis) for 100 parametric simulation runs. See Section Competitor ordination methods for the meaning of the abbreviations. All samples have the same mean taxa composition, counts were sampled from the negative binomial distribution. A small pseudo-F value is preferred. Results from the RC(M) method are shaded in grey. Top: Four groups with differences in library sizes. Bottom: Four groups with differences in dispersions.

**Fig 6.**
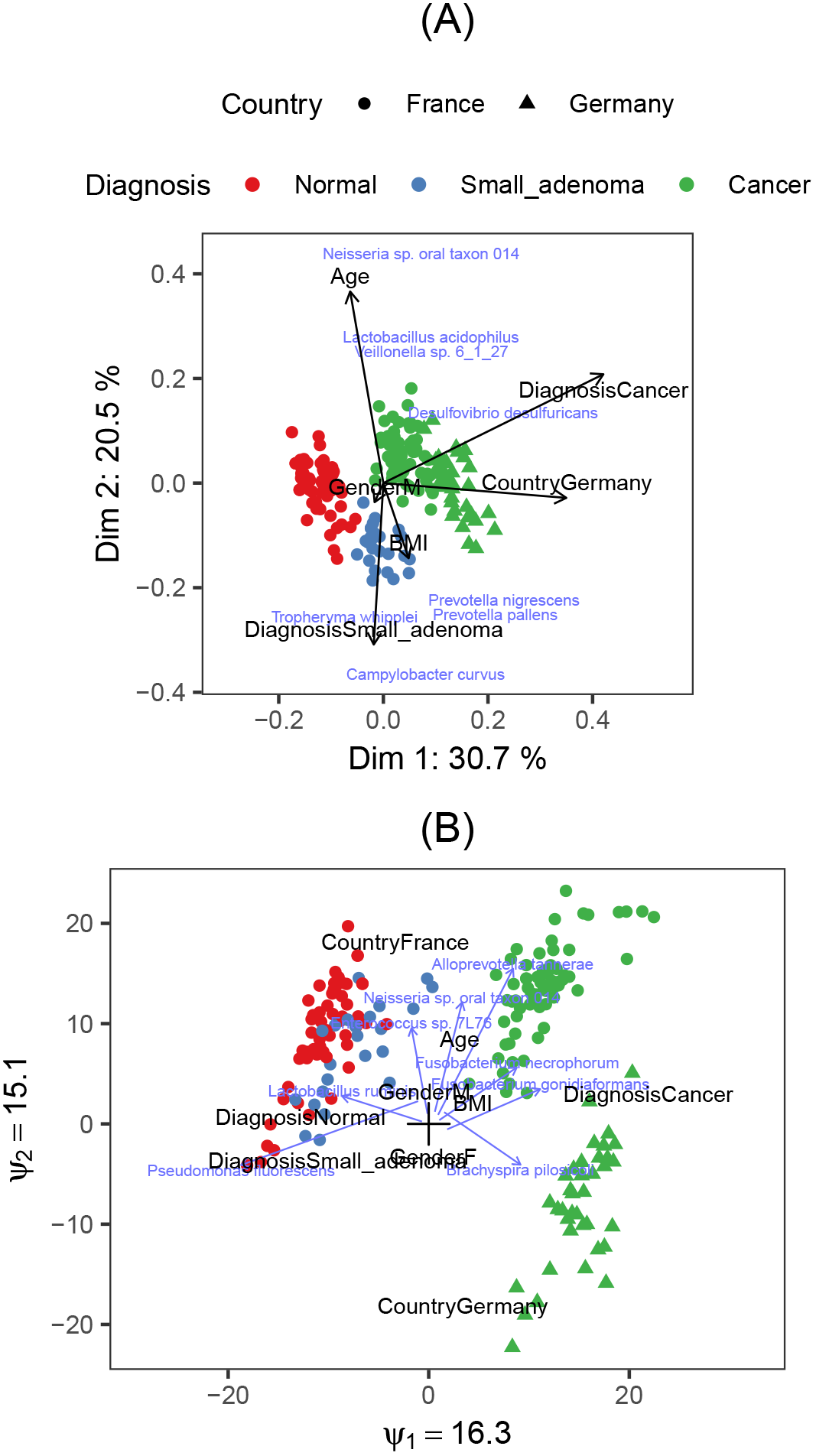
Results of biological signal simulations. Boxplots of the silhouette (top), pseudo-F statistic (center) and taxon ratio (bottom) for several ordination methods (x-axis) over 100 parametric simulation runs. See Section Competitor ordination methods for the meaning of the abbreviations. 10% of the taxa were made differentially abundant in each of 4 sample groups, with a fold change of 5. A large pseudo-F value is preferred. Columns correspond to the simulation scenario: negative binomial (NB) (cor: data generation with taxon correlation, phy: phylogenetically correlated taxa were made differentially abundant), Dirichlet multinomial (DM) and zero-inflated negative binomial (ZINB). Results from the RC(M) method are shaded in grey.

### Simulation study

#### No-Signal Simulations

Fig 5 shows the pseudo F-statistics for the no-signal simulations with the negative binomial distribution and with four groups of different library sizes or different dispersions. Since sequencing depths are assumed to be unrelated to the biological composition of a sample [10, 27], they should not affect the sample ordinations by, for example, forming clusters of samples with similar library sizes. Many methods appear to be insensitive to library size variability, except the ordinations based on Hellinger distances, PCoA with Bray-Curtis dissimilarities on absolute and logged abundances, and the compositional data analysis (CoDa). Their sensitivity to the library sizes can also be seen in S1 Fig, where the correlations between the sample scores and the library sizes for the first three dimensions is shown. It has been noted before that distance-based methods are sensitive to differences in dispersion between different sample groups [3, 12]. Our simulations confirm that all PCoA methods investigated, as well as CoDa, Hellinger distance and our RC(M) method tend to cluster samples with the same dispersion levels together, even when all samples have equal taxa compositions (see Fig 5).

#### Biological Signal Simulations

As shown in Fig 6, he biological signal is best detected with the RC(M) method (large Silhouette and pseudo-F values) and RC(M) succeeds best in identifying the driving taxa (large taxon ratio). This holds for all scenarios, except for data generated by the Dirichlet multinomial (DM) distribution. Also detrended correspondence analysis (DCA) is good at detecting the important taxa. More results, with similar conclusions, can be found in Section 3 of the S1 Appendix.

## Discussion

Unconstrained and constrained ordination techniques that are currently employed in microbial ecology rely mainly on eigenvalues/eigenvectors and singular value decompositions. Although having the advantage of computational efficiency, they are too rigid to deal with some of the more peculiar aspects of microbial amplicon sequencing data. For instance, sequencing depths varying between samples and taxon-wise overdispersions are two characteristics of microbiome data that may distort ordinations. One possible reason why these flaws received little attention, is because their underlying assumptions are rarely stated explicitly, and hence they are almost never checked. Researchers in microbial ecology should become more aware of assumptions and limitations of the ordination methods. Ordination methods developed for ecological data with directly observed species counts may no longer be valid for sequencing data, because sequencing counts are only a proxy of abundance and the biological and technical variability show specific characteristics. Dimension reduction for plotting inevitably entails information loss, but using ordination methods that are inappropriate for the data type may yield misleading results.

Distance-based methods are currently very popular ordination methods in microbiomics. However, by calculating distances between samples, the information on which taxa discriminate the samples is discarded. As a result, distance based methods cannot directly identify which taxa drive the differences between samples, limiting their use for data exploration.

Compositional data analysis (CoDa) analyzes ratios between taxon counts rather than the counts themselves. Although sequencing data often should be treated as compositional indeed, these methods ignore the count origin and the associated heteroscedasticity. As a result, the sample scores of their ordinations correlate strongly with the library sizes, which are considered as technical artefacts. This is highly problematic for the interpretation of their ordination diagrams. Especially in datasets with a low signal-to-noise ratio, differences in library sizes, rather than biological signal, may be depicted in the ordination graphs. Because of the common association of library sizes with sample-specific variables, this may incorrectly confirm the researcher’s prior beliefs in differences in microbiome composition, whereas actually none exist.

Despite their longer computation times, ordination methods based on count regression models are more flexible to deal with these issues, and have gained popularity over the recent years. The RC(M) regression model can include an offset to account for varying sequencing depths, and can be easily augmented with skewed count distributions with taxon-wise parameters to address heteroscedasticity. Furthermore, it can condition out the effect of other confounding variables. The main idea is that interaction terms between samples and taxa capture departures from equal taxa composition in the samples. These interaction terms can then be plotted to visualize the strongest signal in the dataset. These strongest signals need not necessarily come from the most abundant taxa. The likelihood framework in which the RC(M) model is fitted, comes with standard diagnostic tools to assess model assumptions. Moreover, outlying or influential observations can be identified, which can reveal useful information to researchers.

Just as row-column interaction models, correspondence analysis tries to represent departures from sample homogeneity in few dimensions. Still, for skewed and overdispersed data, an additive model for departure from equal sample composition is inappropriate and produces ordination plots dominated by outliers. A multiplicative model as employed in the RC(M) model is more appropriate for these data.

The performance of ordination methods can be assessed quantitatively through simulations. Our comprehensive simulation study confirms a good performance of the RC(M) method, both in terms of sample separation as in the identification of taxa that contribute to these separations. The RC(M) method is not sensitive to library size variation, but, just as many other ordination methods, it is somewhat sensitive to differences in dispersions.

We believe the potential of row-column interaction models is underemployed in the analysis of all types of high-dimensional data, despite the availability of contemporary fitting algorithms and computing power. However, given the reasonably good performance of CoDa techniques in our simulations, a combination of log-linear models that correctly model the mean-variance structure, and models that account for compositionality would probably further improve visualization methods for the microbiome.

Constrained ordinations include sample-specific variables in the visualization. Despite a very rich theoretical foundation, they are less frequently employed in the microbial ecology practice. We combined the row-column interaction model with flexible response modeling using linear response functions as well as non-parametrically estimated response functions. Linear response functions yield easily interpretable triplots, and the linearity assumption can be verified using diagnostic plots. Non-parametrically estimated response function allow maximal flexibility in modelling the taxon niches. Our method uniquely combines unconstrained and constrained ordination into the same framework for fitting and plotting, which greatly facilitates comprehensive exploration of microbiome datasets.

Our methods are implemented in RCM the R-package *RCM* visualization of microbiome data (available at http://github.com/CenterForStatistics-UGent/RCM). The package comes with a custom-written fitting algorithm for the RC(M) model as well as several ready-to-use plotting functions.

## Acknowledgments

Thanks to Ruben Props and Chris Callewaert for fruitful discussions on the application of our method, and to Chris Callewaert for extensively testing the *RCM* R-package.

## Supporting information

**S1 Appendix** A detailed discussion of the RC(M) method, with illustrations on real datasets. Further, a detailed description of the setup and results of the simulation study, followed by a list of software versions.

**S1 File Auxiliary R-code** All R-code for making the graphs shown in the publication, along with the code for the simulation study.

**S2 File Data** All datasets used in this publication.

**S1 Fig Correlations of library sizes and row scores** Boxplots with the correlation of sample scores with observed library sizes (y-axis) for different ordination methods (x-axis). Side panels indicate the different parametric simulation scenarios, see Section Simulation study for an explanation of the codes used. Top panels show the dimension of the sample score

## References

1. Gower JC. In: Principal Coordinates Analysis. John Wiley & Sons, 2005

2. Kruskal JB. Multidimensional scaling by optimizing goodness of fit to a nonmetric hypothesis. Psychometrika. 1964;29(1):1–27.

3. Warton DI, Wright ST, Wang Y. Distance-based multivariate analyses confound location and dispersion effects. Methods in Ecology and Evolution. 2012;3(1):89–101. doi:10.1111/j.2041-210X.2011.00127.x.

4. Weiss S, Xu ZZ, Peddada S, Amir A, Bittinger K, Gonzalez A, et al. Normalization and microbial differential abundance strategies depend upon data characteristics. Microbiome. 2017;5(27). doi:10.1186/s40168-017-0237-y.

5. Wong RG, Wu JR, Gloor GB. Expanding the UniFrac Toolbox. PLOS ONE. 2016;11(9):1–20. doi:10.1371/journal.pone.0161196.

6. Benzecri JP. L’analyse des données. Population. 1975;30(6):1190.

7. ter Braak CJF. Canonical Correspondence Analysis: A New Eigenvector Technique for Multivariate Direct Gradient Analysis. Ecology. 1986;67(5):1167–1179.

8. Zhu M, Hastie T, Walther G. Constrained ordination analysis with flexible response functions. Ecological Modelling. 2005;187:524–536.

9. Zeller G, Tap J, Voigt AY, Sunagawa S, Kultima JR, Costea PI, et al. Potential of fecal microbiota for early-stage detection of colorectal cancer. Mol Syst Biol. 2014;10(766). doi:10.15252/msb.20145645.

10. Gloor GB, Reid G. Compositional analysis: A valid approach to analyze microbiome high-throughput sequencing data. Can J Microbiol. 2016;62(8):692–703. doi:10.1139/cjm-2015-0821.

11. Costea PI, Zeller G, Sunagawa S, Bork P. A fair comparison. Nature Methods. 2014;11:359.

12. Sohn MB, Li H. A GLM-based latent variable ordination method for microbiome samples. Biometrics. 2017; p. e–pub ahead of print. doi:10.1111/biom.12775.

13. Hui FKC, Taskinen S, Pledger S, Foster SD, Warton DI. Model-based approaches to unconstrained ordination. Methods in Ecology and Evolution. 2015;6(4):399–411. doi:10.1111/2041-210X.12236.

14. Anderson MJ, Willis TJ. Canonical analysis of principal coordinates: A useful method of constrained ordination for ecology. Ecology. 2003;84(2):511–525. doi:10.1890/0012-9658(2003)084[0511:CAOPCA]2.0.CO;2.

15. ter Braak CJF, Prentice IC. A Theory of Gradient Analysis. 1988;18(Supplement – C):271–317.

16. Yee TW. Vector Generalized Linear and Additive Models: With an Implementation in R. Springer Series in Statistics. Springer New York; 2015.

17. Goodman L. Simple Models for the Analysis of Association in Cross-Classifications Having Ordered Categories. 1979;74:537–552.

18. Yee T, Hadi A. Row–column interaction models, with an R implementation. 2014;29:1427–1445.

19. Robinson MD, Smyth GK. Moderated statistical tests for assessing differences in tag abundance. Bioinformatics. 2007;23(21):2881–2887. doi:10.1093/bioinformatics/btm453.

20. Yee TW. Constrained additive ordination. Ecology. 2006;87(1):203–213. doi:10.1890/05-0283.

21. R Core Team. R: A Language and Environment for Statistical Computing; 2015. Available from: http://www.R-project.org/.

22. Minchin P. An Evaluation of the Relative Robustness of Techniques for Ecological Ordination. 1987;69:89–107.

23. Faith DP, Minchin P, Belbin L. Compositional dissimilarity as a robust measure of ecological distance. 1987;69:57–68.

24. Legendre P, Gallagher ED. Ecologically meaningful transformations for ordination of species data. Oecologia. 2001;129(2):271–280. doi:10.1007/s004420100716.

25. Kuczynski J, Liu Z, Lozupone C, McDonald D, Fierer N, Knight R. Microbial community resemblance methods differ in their ability to detect biologically relevant patterns. Nat Methods. 2010;7(10):813–819. doi:10.1038/nmeth.1499.

26. Ruokolainen L, Salo K. Differences in performance of four ordination methods on a complex vegetation dataset. Science. 2006;43:269–275.

27. McMurdie PJ, Holmes S. Waste Not, Want Not: Why Rarefying Microbiome Data Is Inadmissible. PLoS Comput Biol. 2014;10(4):e1003531. doi:10.1371/journal.pcbi.1003531.

28. Fukuyama J, McMurdie PJ, Dethlefsen L, Relman DA, Holmes S. Comparisons of distance methods for combining covariates and abundances in microbiome studies. Pac Symp Biocomput. 2012; p. 213–224.

29. Schmidt TSB, Rodrigues JFM, von Mering C. A family of interaction-adjusted indices of community similarity. The Isme Journal. 2016;11(3):791–807.

30. Dray S, Pavoine S, de Cárcer DA. Considering external information to improve the phylogenetic comparison of microbial communities: A new approach based on constrained Double Principal Coordinates Analysis (cDPCoA). Molecular Ecology Resources. 2015;15(2):242–249.

31. Clarke K. Nonparametric Multivariate Analyses of Changes in Community Structure. 1993;18:117–143.

32. Peterson J, Garges S, Giovanni M, McInnes P, Wang L, Schloss JA, et al. The NIH Human Microbiome Project. Genome Res. 2009;19(12):2317–2323. doi:10.1101/gr.096651.109.

33. org A. The American gut project;.

34. Kostic AD, Gevers D, Pedamallu CS, Michaud M, Duke F, Earl AM, et al. Genomic analysis identifies association of Fusobacterium with colorectal carcinoma. Genome Res. 2012;22(2):292–298. doi:10.1101/gr.126573.111.

35. Turnbaugh PJ, Ridaura VK, Faith JJ, Rey FE, Knight R, Gordon JI. The Effect of Diet on the Human Gut Microbiome: A Metagenomic Analysis in Humanized Gnotobiotic Mice. Sci Transl Med. 2009;1(6):6ra14–6ra14. doi:10.1126/scitranslmed.3000322.

36. Props R, Kerckhof FM, Rubbens P, De Vrieze J, Hernandez Sanabria E, Waegeman W, et al. Absolute quantification of microbial taxon abundances. The ISME Journal. 2016;11:584–587.

37. La Rosa PS, Brooks JP, Deych E, Boone EL, Edwards DJ, Wang Q, et al. Hypothesis Testing and Power Calculations for Taxonomic-Based Human Microbiome Data. PLoS ONE. 2012;7(12):e52078. doi:10.1371/journal.pone.0052078.

38. Kurtz ZD, Müller CL, Miraldi ER, Littman DR, Blaser MJ, Bonneau RA. Sparse and Compositionally Robust Inference of Microbial Ecological Networks. PLoS Comput Biol. 2015;11(5):1–25. doi:10.1371/journal.pcbi.1004226.

39. Chen J, Bittinger K, Charlson ES, Hoffmann C, Lewis J, Wu GD, et al. Associating microbiome composition with environmental covariates using generalized UniFrac distances. Bioinformatics. 2012;28(16):2106–2113. doi:10.1093/bioinformatics/bts342.

40. McMurdie PJ, Holmes S. phyloseq: An R package for reproducible interactive analysis and graphics of microbiome census data. PLoS ONE. 2013;8(4):1–11.

41. Gower J, Lubbe S, le Roux N. Understanding Biplots. vol. 1; 2011.

42. Rao CR. A review of canonical coordinates and an alternative to correspondence analysis using Hellinger distance. Qüestiió. 1995;19(1–3):23–63.

43. van der Maaten L, Hinton G. Visualizing data using t-SNE. Journal of Machine Learning Research. 2008;9(Nov):2579–2605.

44. Rousseeuw PJ. Silhouettes: A graphical aid to the interpretation and validation of cluster analysis. Journal of Computational and Applied Mathematics. 1987;20(Supplement – C):53–65. doi:10.1016/0377-0427(87)90125-7.

45. Anderson MJ. A new method for non-parametric multivariate analysis of variance. Austral Ecology. 2001;26(1):32–46. doi:10.1111/j.1442-9993.2001.01070.pp.x.

46. Hiergeist A, Reischl U, Gessner A. Multicenter quality assessment of 16S ribosomal DNA-sequencing for microbiome analyses reveals high inter-center variability. International Journal of Medical Microbiology. 2016;306(5):334–342. doi:10.1016/j.ijmm.2016.03.005.

47. Legendre P, Legendre LFJ. Numerical Ecology. Developments in Environmental Modelling. Elsevier Science; 2012.

48. Macarthur R, Levins R. The Limiting Similarity, Convergence, and Divergence of Coexisting Species. The American Naturalist. 1967;101(921):377–385.

49. van den Wollenberg AL. Redundancy analysis an alternative for canonical correlation analysis. Psychometrika. 1977;42(2):207–219. doi:10.1007/BF02294050.

50. Hastie T, Tibshirani R. Generalized Additive Models. Statistical Science. 1986;1(3):297–310.

51. Yee TW. A new technique for maximum-likelihood canonical gaussian ordination. Ecological Monographs. 2004;74(4):685–701. doi:10.1890/03-0078.

52. Zhang Y, Thas O. Constrained Ordination Analysis with Enrichment of Bell-Shaped Response Functions. PLOS ONE. 2016;11(4):1–21. doi:10.1371/journal.pone.0154079.

53. McCullagh P, Nelder JA. Generalized Linear Models, Second Edition. Chapman & Hall/CRC Monographs on Statistics & Applied Probability. Taylor & Francis;1989.

54. Hampel FR, Ronchetti EM, Rousseeuw PJ, Stahel WA. Robust Statistics: The Approach Based on Influence Functions. vol. 07. John Wiley & Sons, Inc.;2011.

